# Physiologically relevant media are associated with overlapping metabolic responses in primary human hepatocytes and Huh7 cells

**DOI:** 10.64898/2026.03.16.712092

**Authors:** Eloise Cross, Felix Westcott, Kieran Smith, Shilpa R. Nagarajan, Fabio Sanna, Kaitlyn M.J.H. Dennis, Leanne Hodson

## Abstract

**Background:** Metabolic dysfunction-associated steatotic liver disease (MASLD) is challenging to study *in vivo* in humans and in *vitro* models are limited. Although primary human hepatocytes (PHHs) are considered the gold-standard, immortalized hepatic cell lines are utilised due to scalability. This study compared the metabolic responses of PHHs with our Huh7-based model cultured in physiologically-relevant fatty acid (FA) mixtures.

**Methods:** PHH and Huh7 cells were treated with 2% human serum, sugars and FAs enriched in either unsaturated (OPLA) or saturated (POLA) FAs for 4 or 7 days, respectively. Stable isotope tracers investigated basal metabolic changes in response to treatment. Cell viability, media biochemistry, intracellular metabolism, lipid droplet morphology and gene expression were quantified.

**Results:** Huh7 cells had greater viability than PHHs, while NEFA uptake and triglyceride secretion were similar. OPLA and POLA increased large lipid droplets in Huh7 cells, whereas only OPLA produced comparable effects in PHHs. Despite higher baseline TG in PHHs, both models showed similar lipid composition, *de novo* lipogenic responses, and glycogen levels. Compared to Huh7 cells, PHHs exhibited higher 3-hydroxybutyrate, lower lactate, reduced glucose uptake, and donor-dependent transcriptomic variability.

**Conclusions:** Huh7 cells are metabolically adaptable and when cultured in physiologically-relevant media, produce metabolic readouts similar PHH cells.

## Introduction

Metabolic dysfunction-associated steatotic liver disease (MASLD), which encompasses a spectrum of disease states from simple steatosis to steatohepatitis ^[1]^, is initiated with the pathological accumulation of intrahepatocellular triglyceride (IHCTG) ^[2]^. MASLD is estimated to affect approximately 30% of the global adult population ^[3]^ and is an independent risk factor for cardiovascular disease ^[4]^. Although it has been suggested that IHCTG accumulation starts to occur when there is an imbalance in hepatocyte fatty acid (FA) uptake, synthesis (*de novo* lipogenesis (DNL)) and disposal (i.e. as TG in VLDL or utilized in mitochondrial oxidation) ^[5–8]^, studying these processes *in vivo* in humans is challenging as high-fidelity disease models are sparse.

The development and progression of MASLD in humans has been investigated through the use of liver biopsies, which offer valuable histological insights, but are invasive, can be difficult to obtain, limited to a single time point, and do not capture the temporal dynamics or spatial heterogeneity of disease progression. Although preclinical rodent models have been utilized to ascertain various aspects of the MASLD pathophysiology ^[9, 10]^, findings from Ding and colleagues ^[11]^ highlighted key divergences in the hepatic handling of FAs between humans and rodents, which may limit the translation of findings ^[12]^. A variety of *in vitro* human cellular models have been utilized to gain insight into the mechanisms underpinning MASLD initiation and progression *in vivo* in humans, however the usefulness of these models is questionable as their ability to recapitulate human disease is often limited ^[12]^.

Primary human hepatocytes (PHH) and immortalized hepatic cell lines have been used as *in vitro* models of disease. PHH are often considered to be the gold-standard ^[3, 12, 13]^ as observations in PHH cells have not often been recapitulated in immortalized hepatocellular carcinoma cell-lines, such as Huh7 and HepG2 ^[14]^. However, PHH also comes with challenges such as high costs, limited culturing capability, potential for low cell viability and significant variation between donor phenotype and genotype ^[15]^, making them somewhat burdensome for regular use when undertaking new methodologies and techniques or larger scale mechanistic experiments. The divergence in observations between PHH and cell-lines may be due to non-physiological media compositions and culturing processes, with *in vitro* cell-lines studies typically utilizing acute exposures (6-48 hours) to supraphysiological concentrations of glucose, a single FA and/or other substrate treatments ^[12]^. We ^[16]^ and others ^[17–19]^ have previously reported that culturing human cell-lines with human serum (HS), rather than foetal bovine serum (FBS), can alter the phenotype of cell-lines to reflect improved functionality of IHCTG accumulation. Moreover, by exposing cell-lines chronically (7 days of culturing) to more physiologically relevant substrate concentrations of FAs, sugars and HS, our model displayed metabolic characteristics of early stage IHCTG accumulation ^[20, 21]^. It remains unclear whether PHHs recapitulate these early-stage characteristics when cultured in similar media conditions. Therefore, the aim of this study was to compare the metabolic responses of PHHs with our previously characterized Huh7-based model ^[20]^ when exposed to the physiologically relevant FA mixtures, OPLA (predominantly unsaturated) and POLA (predominantly saturated), as a model of early IHCTG accumulation. Given the differing viability and culture requirements of PHH and Huh7 cells, we compared each model under its optimal chronic culture conditions, rather than attempting perfectly matched exposure duration or basal media composition.

## METHODS

### Cell Culture

Huh7 and PHH cells were used. Huh7 cells were kindly provided by Dr Camilla Pramfalk (Karolinska Institute) and were cultured as previously described ^[20, 21]^. PHH cells were obtained from Thermofisher (ThermoFisher, HMCPSQ) originating from both male and female donors (see supplementary data, Table 1 for donor characteristics. Frozen PHH cells were thawed in a 37°C water bath rapidly. 500µL of warm plating media (500mL Williams E Medium (ThermoFisher, A1217601) supplemented with a Primary Hepatocyte Thawing and Plating Supplements (ThermoFisher, CM3000) which contained FBS, dexamethasone, and a cocktail of penicillin-streptomycin, insulin, GlutaMAX™ and HEPES solution, was then added to the cryovial. Cells were then suspended and transferred to a falcon tube containing 4.5mL of warm plating media. Cell number and viability was then quantified using Trypan blue stain (ThermoFisher, 15250061) and a Cellometer Auto T4 Bright Field Cell Counter. Cells were then diluted with plating media and seeded into plates which had been pre-coated with collagen type I (ThermoFisher, A1048301) at a density of 35,000 cells/well for 96-well plates, 150,000 cells/well for 24-well plates or 50,000 cells/well for 12mm diameter glass cover slips for microscopy experiments. Both Huh7 and PHH cells were maintained at 37°C in 5% CO_2_.

### Experimental Treatments

Huh7 cells were seeded into relevant plates and cultured in experimental media comprised of 2% HS, 5.5mM glucose and 800µM of 4 FAs (oleic acid (O), palmitic acid (P), linoleic acid (L) and alpha-linoleic acid (A)) in one of two physiological ratios: OPLA (45:30:24:1) or POLA (44:45:10:1), or a no fat control as previously described ^[20]^. Huh7 cells were cultured in experimental media for seven days with media changes every other day as previously described ^[20]^.

Following seeding, PHHs were cultured in plating media overnight then cultured in experimental media which consisted of Williams E Medium (ThermoFisher, A1217601) and Primary Hepatocyte Maintenance Supplements (ThermoFisher, CM4000) containing dexamethasone and a cocktail of penicillin-streptomycin, insulin, transferrin, selenium complex, BSA, GlutaMAX™, and HEPES solution, supplemented with 2% HS. PHH cells were cultured in 800µM OPLA and POLA or no fat control experimental media for four days with media changes every other day.

### Biochemical Analysis

Media glucose, lactate, TG, apolipoprotein B (apoB0) and non-esterified FA (NEFA) concentrations were measured on a AU480 chemistry analyzer (Beckman Coulter) and normalized to protein concentration as described ^[16]^. Media ATP (Promega), acetate (Abcam) and intracellular glycogen (BioVision) were measured using plate-reader assays according to manufacturer’s instructions.

### Cell viability

We assessed cell viability in two ways. Prior to seeding in experimental media, Trypan Blue, which stains cells with an intact plasma membrane, was added to Huh7 and PHH cells. Cellular ATP concentrations were determined using the CellTiter[Glo Luminescent Cell Viability Assay (Promega; WI, USA) according to manufacturer’s instructions. Luminescence was read on FLUOstar Omega (BMG Labtech). For this, we also assessed the effect of individual FAs and FA mixtures (i.e., OPLA and POLA) or a no-fat control, on cell viability by culturing Huh7 and PHH cells in media supplemented with 400µM oleate or 400µM palmitate as either a single FA or as part of a FA mixture (800µM OPLA or 800µM POLA) with the media changed daily for 4 days.

### Lipid Extraction and Gas Chromatography-Mass Spectrometry

To assess intrahepatocellular DNL, deuterium (10% v/v) was added to experimental media for the entire duration of the treatment (i.e. either 2, 4, or 7 days). Cells were harvested and the TG and phospholipid fractions and quantified and DNL measured as described ^[20]^.

### Confocal Microscopy

To assess lipid droplet (LD) size, Huh7 and PHH cells were seeded on 12mm diameter glass cover slips (VWR, 631-1577) in 24-well plates. After the respective treatment period, cells were washed with PBS and then fixed by incubation with 4% paraformaldehyde in PBS for 15mins. Cover slips were stored in PBS at 4°C until staining. To allow penetration of the stains into cells, cover slips were kept in 0.01% Triton-X in PBS for 25mins. Coverslips were then washed twice with PBS and stained with HCS LipidTOX™ Red Neutral Lipid Stain (ThermoFisher, H34476) at 1:200 in PBS and RedDot™2 Far-Red Nuclear Stain (Biotium, 40061) at 1:125 in PBS for 1 hour followed by two more washes with PBS and staining with Oregon Green™ 488 Phalloidin (ThermoFisher, O7466) at 1:40 in PBS for 30mins. Coverslips were then washed twice with PBS and mounted onto a slide with VECTASHIELD Vibrance® Antifade Mounting Medium (Vector Laboratories, H-1700) before imaging with a confocal microscope (Bio-Rad, Radiance 2100). Images were analysed using open source CellProfiler4 software.

### RNA extraction

RNA was extracted using the RNEasy^®^ Plus mini kit (QIAGEN Sciences) according to manufacturer’s instructions. The resulting RNA was treated with DNase (QIAGEN Sciences) and quality checked with an Agilent Bioanalyzer.

### RNA sequencing analysis

Purified RNA samples from 5 donors (see supplementary data, Table 1 for donor characteristics) were sent to Novogene for bulk RNA sequencing. Novogene carried out the initial data processing which involved the following: Removal of low-quality reads or reads containing poly-N regions using in-house perl scripts, indexing and alignment of clean reads to a downloaded reference genome using Hisat2 (version 2.0.5) and finally counting reads mapped to each gene using featureCounts (version 1.5.0-p3). The following was carried out on all datasets in R (version 4.4.1) on RStudio (version 2024.04.2+764). Raw gene count data was filtered to remove genes with low expression, and then normalised using the TMM method ^[22]^ with the EdgeR package (version 4.2.1) ^[23]^. FactoMineR and factoextra packages were used to generate principal component analysis (PCA) plots. Differential gene expression analysis using a mixed effects model to control for differences between donors/replicates was performed using the limma package (version 3.60.4) to remove heteroscedasticity from the data ^[24]^. Genes were considered to be differential expressed if the false discovery rate (FDR) <0.05. To determine which pathways were differentially regulated between conditions, gene set enrichment analysis (GSEA) was performed with the package clusterProfiler (version 3.0.4) using the Molecular Signatures Database (MSigDB) Hallmark gene set collection ^[25]^.

### Statistical Analysis

All experiments were carried out with technical duplicates with at least three independently performed biological repeats. Data are presented as mean±standard deviation (SD) unless otherwise specified. For comparisons involving two independent groups, the Mann–Whitney U test was used. When comparing multiple groups, the Kruskal–Wallis test followed by Dunn’s multiple comparisons test was used. Statistical analyses were conducted using GraphPad Prism (version 10). A p-value of <0.05 was considered statistically significant.

## RESULTS

PHH and Huh7 cells were exposed to basal medias and supplements that differed in composition and cells were cultured in treatment media for a different amount of time, 7 days for Huh7 cells and 4 days for PHH cells, which may have influenced the viability and metabolism of cells.

### Cell viability

When cell viability was assessed, prior to seeding in experimental media, with Trypan Blue, we found 95.6±4.5% of Huh7 cells were viable, whereas only 68.0±9.3% of PHH cells were viable across 5 seeding batches suggesting that prior to seeding PHH cells have lower viability than Huh7 cells.

Although we found no significant differences in ATP levels across conditions in both Huh7 and PHH cells, the PHH cells had significantly lower ATP levels across all conditions (Figure 1A-B). There was a higher degree of variability in ATP concentrations of Huh7 cells cultured in palmitate alone compared to other conditions, which was not observed in PHH cells (Figure 1A-B). Our results indicate that Huh7 cells have increased viability compared with PHH cells, and the different cell types may exhibit differing metabolic response in response to culturing with only palmitate.

**Figure 1:**
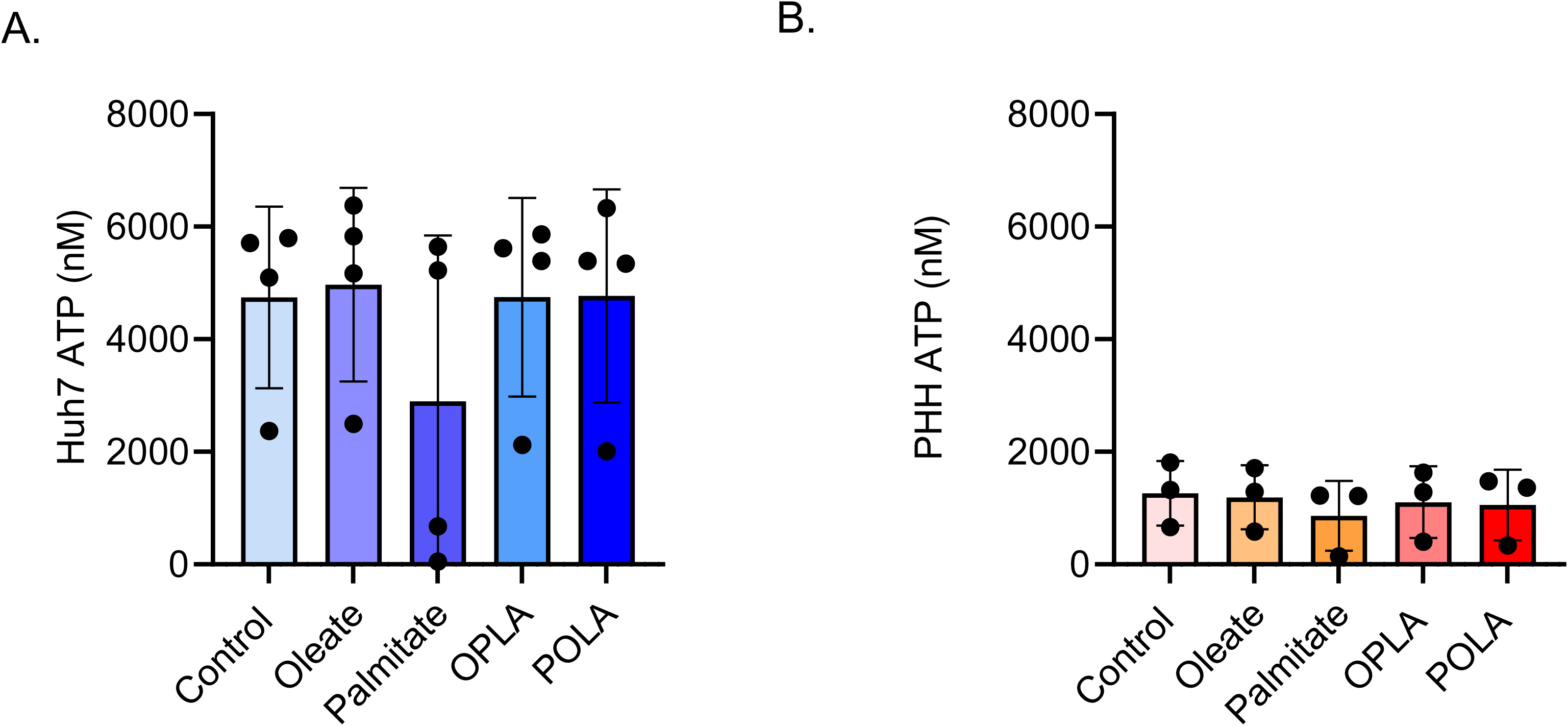
The effects of a single FA and FA mixtures enriched in unsaturated or saturated fatty acids on cell viability. Treatments consisted either of control (no FAs; 5.5 mM glucose), palmitate (400 µM palmitate; 5.5 mM glucose), oleate (400 µM oleate; 5.5 mM glucose), OPLA (800 µM FAs; 5.5 mM glucose) or POLA (800 µM; 5.5 mM glucose) media where FA mixtures were enriched in either unsaturated (O) or saturated (P) FAs. Intracellular ATP was measured as a marker of cell viability in **A)** Huh7 (n = 4) and **B)** Primary Human Hepatocytes (PHH) (n=3) cells. Data are presented as ± SD using Kruskal–Wallis test followed by Dunn’s multiple comparisons post-hoc test for A-B.

### Fatty acid uptake and Media TG

Uptake of media NEFA from OPLA and POLA was assessed in Huh7 and PHH cells. In both OPLA- and POLA-treated Huh7 cells, approximately 80% of media NEFA was taken up (Figure 2A). In contrast, in both OPLA- and POLA-treated PHH cells media NEFA uptake was approximately 70% (Figure 2B). Although NEFA uptake was broadly similar between models, greater variability was observed in PHH, consistent with donor heterogeneity (Supplementary Table 1). Occasional lower uptake values were observed across experiments, highlighting inhered biological and technical variability in PHH cultures. For both Huh7 and PHH cells, there were no significant differences in media TG concentrations between treatments, although there was greater variation within the FA treatments in PHH than Huh7 cells (Figure 2C-D).

**Figure 2:**
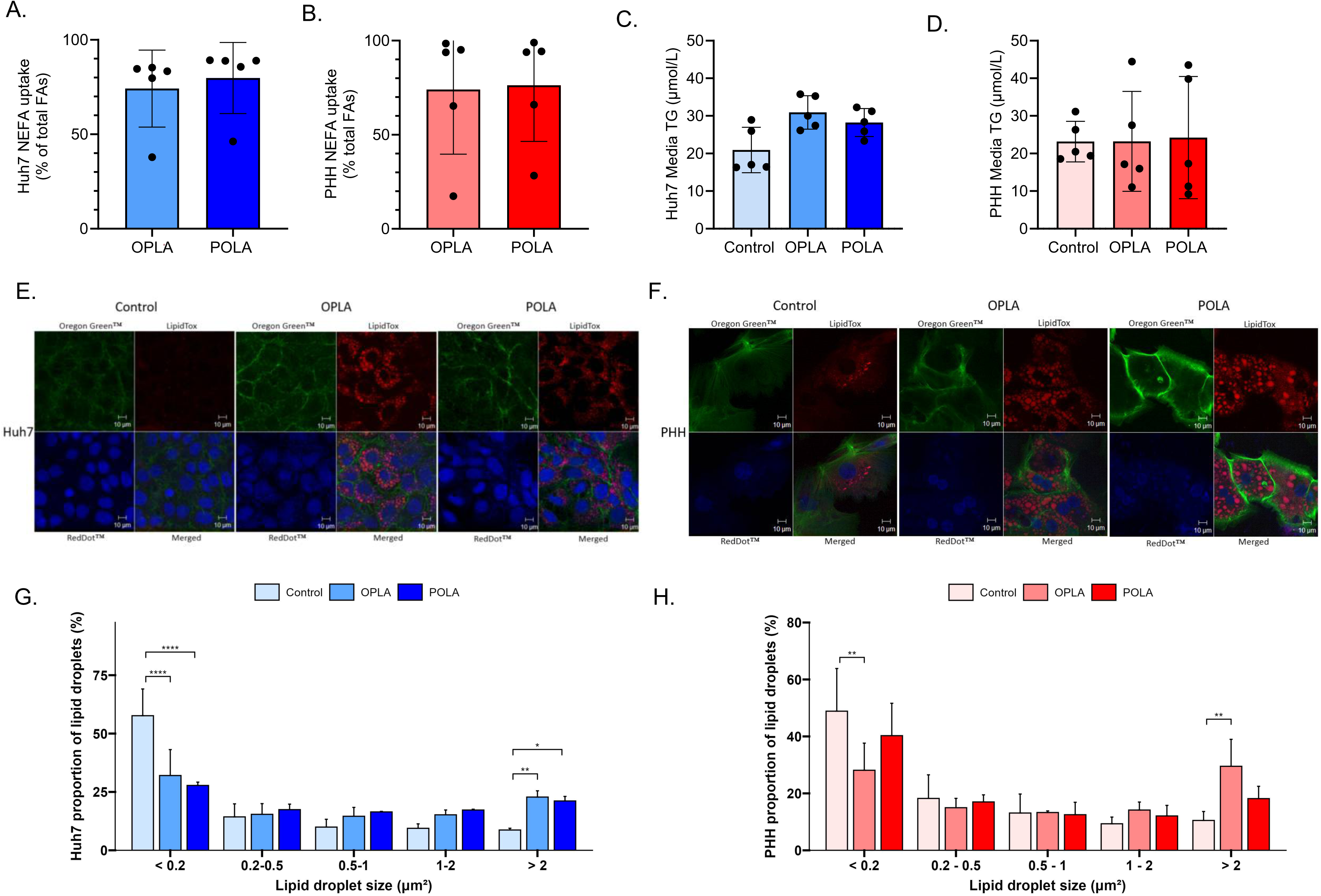
The effects of fatty acid mixtures on fatty acid uptake, secretion and lipid drop size in Huh7 and PHH cells. Non-esterified fatty acid (NEFA) uptake from the media in **A)** Huh7 and **B)** PHH cells, media triglyceride (TG) concentrations **C)** Huh7 and **D)** PHH cells and lipid droplet imagining **E)** Huh7 and F) PHH cells, shown in representative confocal microscopy images taken at 60x magnification of cells cultured in no-fat control, 800 µM OPLA or 800 µM POLA and the proportion of lipid droplets across being assess from the image analysis from at least six images per biological repeat allowed profiling of the lipid droplet size in **G)** Huh7 and **H)** PHH cells. Data are presented as ± SD *p<0.05, **p<0.01, ****p<0.0001 (Mann-Whitney U test for A-D; two-way ANOVA followed by Tukey’s HSD multiple comparisons post-hoc test for G-H).

### Lipid droplet size

The formation of LDs was observed in both Huh7 and PHH cells and appeared greater in OPLA- and POLA-treated cells compared with cells treated with no-fat control media (Figure 2E-F). These observations were confirmed following image quantification, with both OPLA- and POLA-treated Huh7 cells displaying a larger proportion of LDs >2µm^2^ and a smaller proportion of LDs <0.2µm^2^ compared to no-fat control media treated cells (Figure 2G). In contrast, OPLA-treated PHH cells had significantly more LDs >2µm^2^ and fewer LDs <0.2µm^2^ than PHH cells cultured in no-fat control media, whereas POLA-treated PHH cells did not (Figure 2H). There were no significant differences in LD size distribution between OPLA- and POLA-treated Huh7 or PHH cells (Figure 2G-H).

### Intrahepatocellular TG accumulation

In both Huh7 and PHH cells IHCTG content significantly increased with OPLA-treatment but only tended (P=0.058) to increase with POLA-treatment compared with no-fat control treated cells (Figure 3A-B). No-fat control treated Huh7 cells had minimal IHCTG accumulation compared to PHH cells which had an IHCTG concentration over 20 times more (Figure 3A-B) suggesting PHH cells started with a notably higher IHCTG content than Huh7 cells. This higher baseline lipid burden suggests PHH may retain donor-specific steatotic characteristics, which could potentially constrain further lipid storage responses *in-vitro*.

**Figure 3:**
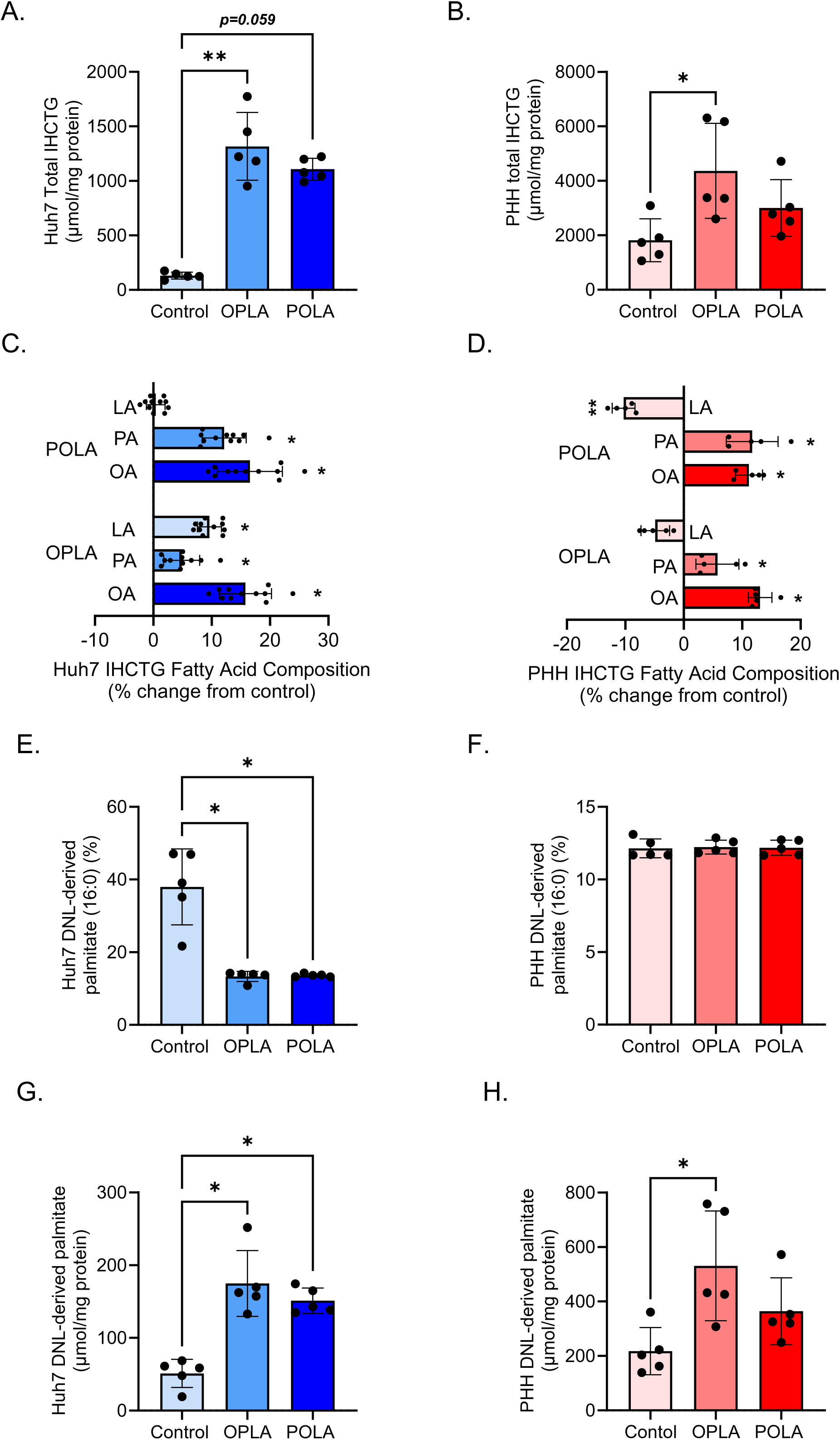
The effects of fatty acid mixtures on intrahepatocellular accumulation and DNL in Huh7 and PHH cells. The intrahepatocellular triglyceride (IHCTG) content of **A)** Huh7 and **B)** PHH cells following culturing in no-fat control, OPLA and POLA treatments and the reflected change in the IHCTG fatty acid composition in **C)** Huh7 and **D)** PHH cells after treatment. The proportion (%) of DNL-derived palmitate was assessed in **E)** Huh7 and **F)** PHH cells along with the absolute contribution of DNL-derived palmitate to IHCTG in **G)** Huh7 and **H)** PHH cells. Data are presented as ± SD *p<0.05, **p<0.01 (Kruskal–Wallis test followed by Dunn’s multiple comparisons post-hoc test for A-B, E-H; Mann-Whitney U test for C-D).

We characterized the FA composition of the accumulated IHCTG content in both Huh7 and PHH cells. We found changes in the IHCTG composition of OPLA- and POLA-treated Huh7 cells to reflect the respective FA media composition when compared to the no-fat control cells (Figure 3C). When compared to no-fat control cells, the proportions of oleate, palmitate and linoleate were significantly higher in OPLA-treated cells, whilst there was no difference between linoleate levels in the no-fat control and POLA-treated cells (Figure 3C). Similarly, for PHH cells, those treated with OPLA and POLA had increases in proportions of oleate and palmitate, reflective of the media compositions (Figure 3D). In contrast, linoleate levels in PHH POLA-decreased significant, whereas there was no notable change in OPLA-treated cells compared to the no-fat control cells (Figure 3D).

### DNL

We found the percentage contribution of DNL-derived palmitate to total IHCTG was significantly lower, at around 13% in OPLA- and POLA-treated Huh7 cells compared to no-fat control cells which were around 38% (Figure 3E). In PHH cells the percentage contribution of DNL-derived palmitate was similar, at 12%, across all treatments. When we accounted for pool size, we found the amount of DNL-derived palmitate in IHCTG to be significantly lower in no-fat control compared to OPLA- and POLA-treated Huh7 cells (Figure 3G), reflecting the greater amount of IHCTG with FA treatment. In PHH cells, although OPLA-treated cells had a significantly higher contribution of DNL-derived palmitate, POLA-treated cells did not, when compared to no-fat control cells (Figure 3H). Although we did not directly compare the contribution of DNL-derived palmitate to IHCTG between FA treated Huh7 and PHH cells, there was a higher amount in PHH (between 249 - 758 µmol/mg protein) compared to Huh7 (between 132 - 251 µmol/mg protein), which likely reflects the higher IHCTG that PHH cells started with.

### Glucose uptake, storage and oxidation

For Huh7 cells cultured in 5mM of glucose, there were no significant differences observed between conditions in the uptake of media glucose between media changes (Figure 4A). PHH cells were cultured in 11mM of glucose, and while there were no significant differences in the absolute glucose uptake between PHH control, OPLA- and POLA-treated cells (Figure 4B), approximately 40% of the available media glucose remained in the media at the end of the culturing period. In contrast, only ∼5% of media glucose remained in the Huh7 cell media, consistent with a greater overall utilization of available glucose under these conditions.

**Figure 4:**
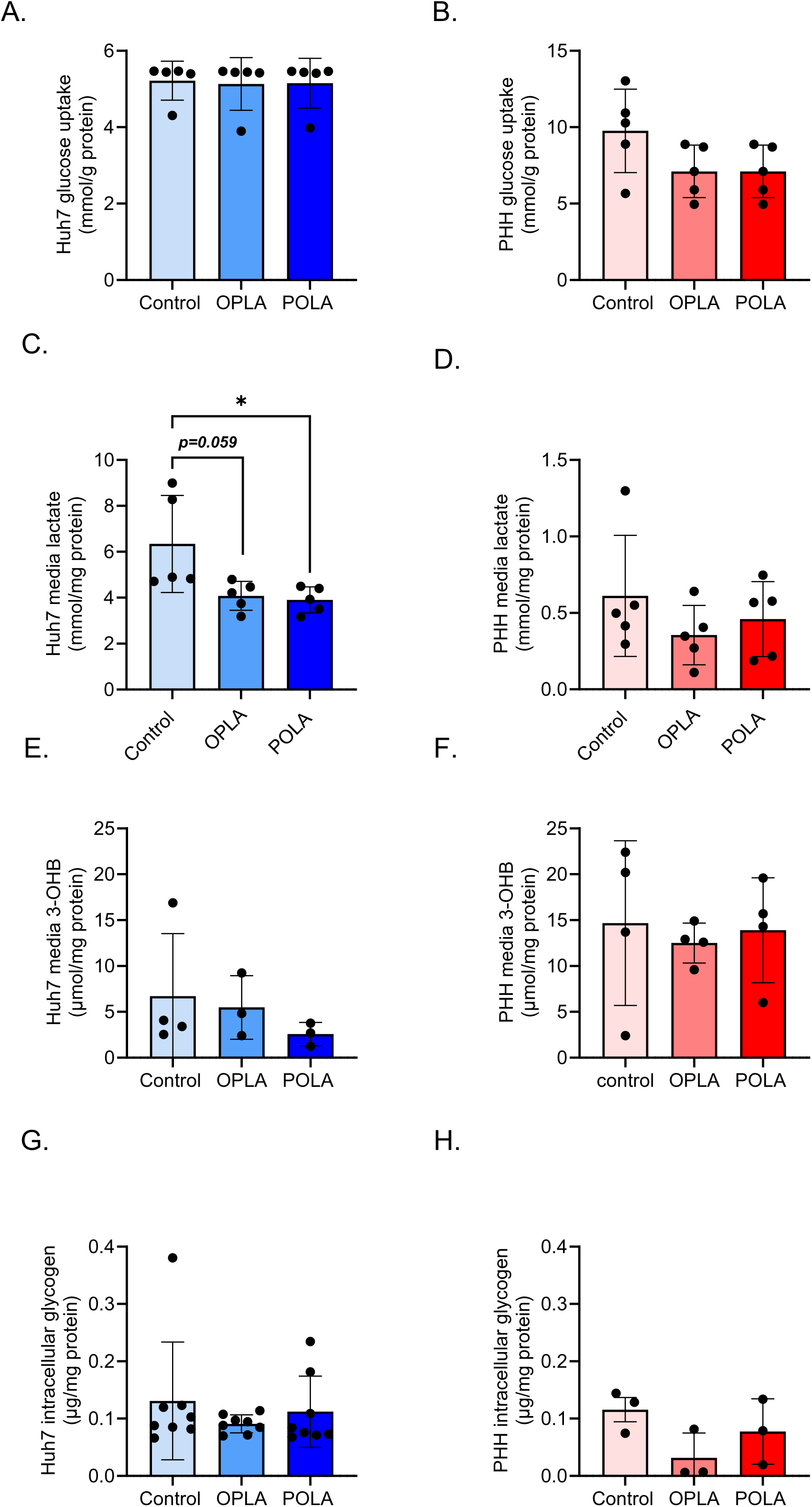
The effects of fatty acid mixtures on glucose metabolism in Huh7 and PHH cells. **A)** Glucose uptake, **C)** Media Lactate, **E)** Media 3-hydroxybutryate (3-OHB) and **G)** intrahepatocellular glycogen concentrations in Huh7 cells, and **B)** Glucose uptake, **D)** Media Lactate, **F)** Media 3-hydroxybutryate (3-OHB) and **H)** intrahepatocellular glycogen concentrations in PHH cells. Data are presented as ± SD using Kruskal–Wallis test followed by Dunn’s multiple comparisons post-hoc test for A-H.

Media lactate levels tended (p=0.058) to be lower and were significantly lower in OPLA- and POLA-treated compared to no-fat control-treated Huh7 cells, respectively (Figure 4C). In contrast, there were no differences in media lactate between conditions in PHH cells (Figure 4D), which were notably lower than the media lactate levels from Huh7 cells.

Although there were not statistically significant differences in 3-hydroxybutyrate (3-OHB) in both Huh7 and PHH cells treated with OPLA, POLA and no-fat control media (Figure 4E-F), media concentrations appeared to be higher in PHH compared to Huh7 cells. No significant differences were observed in intracellular glycogen concentrations in both Huh7 and PHH no-fat control, OPLA-and POLA-treated cells (Figure 4G-H).

### RNA Seq data

Principal component analysis of Huh7 cell transcriptomic data showed the difference in gene expression between Control and OPLA/POLA conditions accounted for the largest component of the total variance (dimension 1 – 65.2%, Figure 5A) in the data and the difference in gene expression between OPLA and POLA conditions accounted for the second largest component (dimension 2 - 9.2%). Gene set enrichment analysis (GSEA) showed that proliferation, oxidative phosphorylation and FA metabolism pathways were the most significantly activated whereas inflammation, unfolded protein response and mammalian target of rapamycin complex 1 (MTORC1) pathways were the most significantly suppressed pathways in both OPLA- and POLA-treated Huh7 cells compared to no-fat control-treated Huh7 cells (Figure 5B-C). When comparing Huh7 cells treated with OPLA and POLA, pathways associated with proliferation were upregulated with POLA treatment and complement, coagulation and cholesterol homeostasis pathways were upregulated with OPLA treatment (Supplementary Figure 1A).

**Figure 5:**
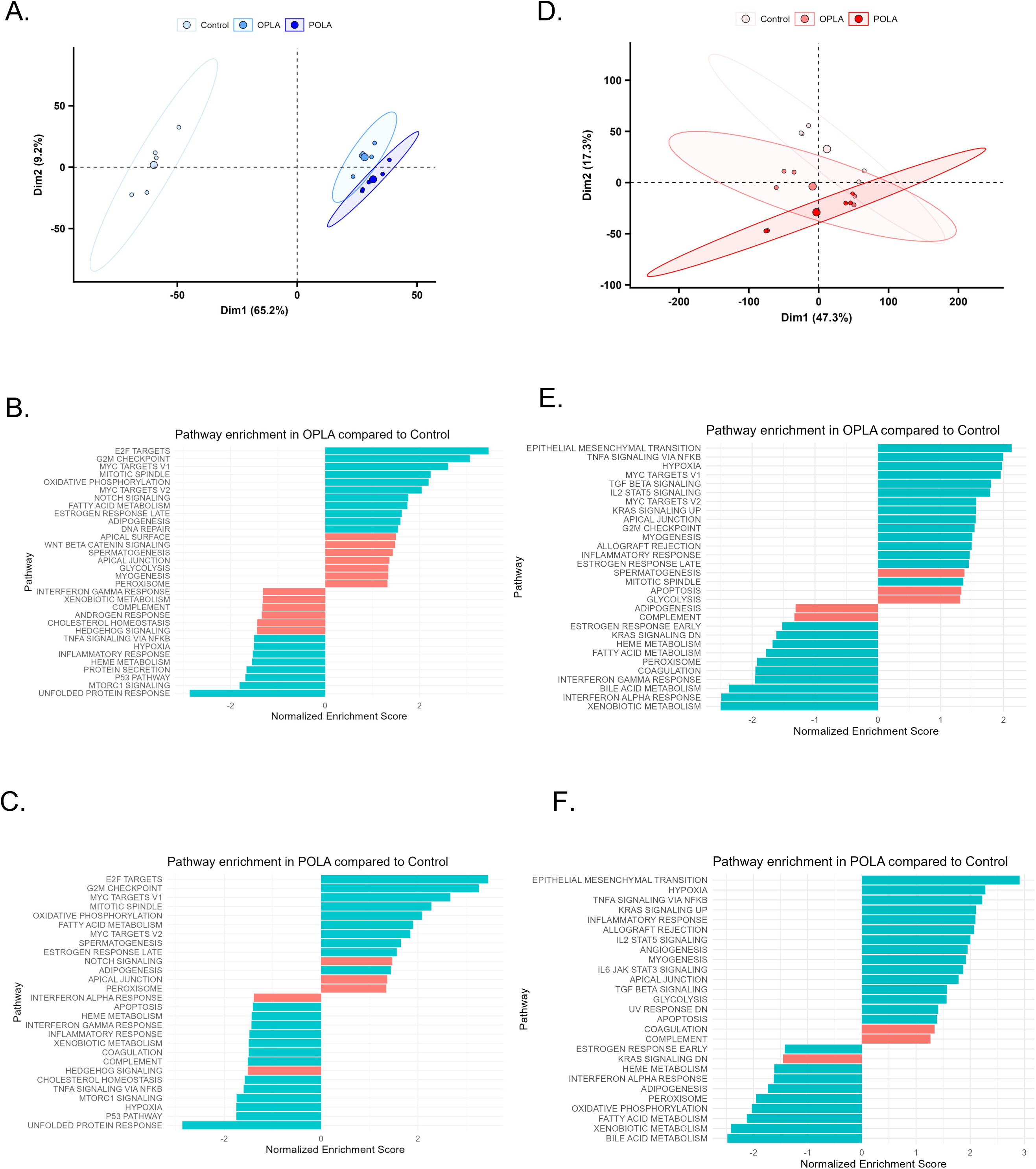
The effects of FA mixtures on gene expression in Huh7 and PHH cells. Principal component analysis plots showing the first two dimensions (dim1 and dim2) for **A)** Huh7 and **B)** PHHs cells treated with no-fat control, OPLA or POLA. Gene set enrichment analysis for OPLA-treated compared to Control-treated cells in **C)** Huh7 and **D)** PHH cells and for POLA-treated compared to Control-treated in **E)** Huh7 and **F)** PHH cells. Pathways with the highest and lowest normalised enrichment score are shown with blue pathways p<0.05 and red pathways p≥0.05 (GSEA performed with FDR correction for multiple comparisons).

In contrast, the largest component of total variance in the PHH transcriptomic dataset was driven by a difference in gene expression between donors (dimension 1 – 47.3%, Figure 5D). However, the second largest component of total variance appeared to be the difference in gene expression between no-fat control and OPLA/POLA conditions (dimension 2 - 17.3%). GSEA highlighted the activation of inflammatory pathways and epithelial mesenchymal transition and the suppression of FA and bile acid metabolism in both OPLA and POLA conditions when compared to no-fat control (Figure 5E-F). Comparison between PHH cells treated with OPLA and POLA showed inflammatory and coagulation pathways upregulated with POLA treatment and proliferation, oxidative phosphorylation and FA metabolism pathways upregulated with OPLA treatment (Supplementary Figure 1B). These findings indicate that, in contrast to Huh7 cells, PHH transcriptional responses to FA exposure are strongly influenced by donor-specific baseline differences.

## DISCUSSION

There is currently a lack of high-fidelity models, particularly *in vitro* cellular models, that recapitulate FA and glucose metabolism, along with IHCTG accumulation observed in early stages of human MASLD. Although PHH are considered to be the gold standard for *in vitro* hepatocellular models, hepatoma cell-lines (i.e. Huh7 and HepG2) appear to be used more routinely as an experimental model. Data comparing PHH and cell-lines as a model to interrogate MASLD initiation and progression is sparse, therefore we compared the metabolic responses of PHHs with our Huh7-cell model of IHCTG accumulation ^[20]^ using chronic (4-7 days) of exposure to physiologically relevant substrates. Despite PHH starting with a higher amount of IHCTG than Huh7 cells, we observed similarities in key metabolic responses including NEFA uptake and IHCTG FA composition, media TG concentrations (reflecting TG secretion), LD morphology, DNL responses, and glycogen levels. Notable differences in responses included PHH having lower and more variable viability than Huh7 cells, higher media 3OHB and lower media lactate concentrations, along with divergence in transcriptomic datasets which, for PHH cells was mostly driven by the gene expression between donors. Together, these data show that Huh7 cells cultured in physiologically relevant media reproduce key lipid-handling phenotypes observed in PHH cells, supporting their use as a scalable workhorse model for mechanistic studies of early IHCTG accumulation. However, PHH remain essential for capturing donor-depended variability and inflammatory stress responses relevant to MASLD progression.

IHCTG accumulation is suggested to be due to an imbalance between input, synthesis and disposal pathways ^[26]^. It is well established that exposing hepatocytes to increasing concentrations of dietary FAs *in vivo* ^[27–30]^ or media FAs *in vitro* ^[20, 21, 31]^ can lead to an increase in IHCTG accumulation. In line with previous *in vitro* studies that have tended to culture hepatocytes short-term (<48h) in a single FA or mix of FA ^[31–35]^, we found both PHH and Huh7 cells accumulated IHCTG with FA treatment compared to the no-fat control. The increase in IHCTG was more evident in Huh7 than PHH cells, despite no notable differences in FA uptake or the FA composition of IHCTG between cell types as both PHH and Huh7 reflected the respective media FA composition after treatment. The lower accumulation of IHCTG by PHH cells, may in part, be explained by the cells having approximately 4 times more IHCTG than Huh7 cells, suggesting a more limited capacity for IHCTG storage. We assessed the change in IHCTG FA composition, compared to the no-fat control cells and found for both PHH and Huh7, it reflected the composition of the FA-treatment. There was a notable decrease in linoleate in POLA-treated PHH cells, which is likely due to the dietary fat the PHH donors were exposed to, in the days prior to hepatocyte isolations, being turned over and replaced with the FA in the culture media, where the linoleate content of POLA was only 10%.

Previous work has shown that Huh7 cells cultured with palmitic acid accumulate less IHCTG than those cultured with oleic acid ^[36–41]^, which may be due to oleic acid promoting storage of FAs in LDs ^[42]^ and palmitic acid activating pro-apoptotic pathways ^[32–35]^. Work by Teixeira and colleagues ^[43]^ demonstrated when HepG2 cells are cultured in 1mM of palmitic acid, linoleic acid or a 1:1 ratio of the two, IHCTG content was highest when cells were cultured with linoleic acid alone, or in combination with palmitic acid. As we wanted to make our media physiologically-relevant, we chose to utilise a mixture of FAs that reflect the major dietary FAs and those found in the human body ^[44]^ and manipulated the proportions of saturated and polyunsaturated FAs. We compared our FA treatments to the more historically used single FA treatments of palmitate and oleate and found that palmitate treatment alone resulted in higher variability in ATP levels in Huh7 cells, a pattern not observed in PHHs. Our findings suggest that even though PHH had lower viability than Huh7, the use of mixed FA chronically (4-7 days) reduced any confounding effects associated with acute lipotoxicity ^[32, 35]^. Thus, media composition can have major effects on the validity, reproducibility and physiological relevance of the scientific findings, therefore choosing an appropriate cell culture medium matters, particularly in metabolic studies ^[45]^.

Although we found PHH cells to have a higher amount of IHCTG than Huh7 cells, we did not observe notable differences in proportions of LD sizes across the two cell types. In Huh7 cells, IHCTG content increased with FA treatments, as did the LD size, whilst in PHH cells only culturing in OPLA, compared to POLA induced a significant increase in IHCTG content and the proportion of larger LDs. Kwon et al ^[13]^ cultured PHH cells in a 3D collagen sandwich for up to 7 days with a FA mix of palmitic and oleic acids (ratio 1:5) in varying concentrations and noted an increase in LD size with higher concentrations of FAs. Paramitha and colleagues ^[46]^ used Raman spectroscopy to compare the effects of specific FA on LD size in HepG2 cells, found the average LD size increased between day 1 and 5 by 11%, 41% and 78% in cells cultured in media, which was changed every 24h, containing 100µmol of palmitic, oleic or linoleic acid, while average LD number per cell increased by 58%, 599% and 233%, respectively. Although we did not assess the number of LDs, it is plausible there was a greater increase in the number of LDs with IHCTG accumulation in Huh7 compared to PHH cells, where there was likely a more notably change in LD number in cells cultured in unsaturated compared to saturated FAs. As LDs are complex, metabolically active organelles in hepatocytes, a notable increase in number and size may cause hepatocyte ballooning and at a whole organ level, lead to anatomical changes in the liver ^[47]^ which may lead to metabolic dysregulation at the whole-body level. Thus, understanding the mechanisms governing LD number and size, along with the impact FA composition may have, will add insight for the further development of therapeutic strategies.

An upregulation in hepatic DNL is a hallmark of insulin resistance and MASLD and has been suggested to promote IHCTG accumulation ^[48]^. We found the percentage contribution of DNL-derived palmitate to IHCTG palmitate to be similar in both cell types with FA-treatment and comparable with what we have previously reported for fasting DNL, *in vivo* in metabolically-healthy humans ^[49–52]^. For the no-fat control treated cells, there was a higher percentage contribution in Huh7 compared to PHH cells, with the latter being similar to the FA-treated PHH and Huh7 cells. After accounting for differences in IHCTG palmitate pool size between the cells, the contribution of DNL-derived palmitate was greater in PHH, reflective of the larger amount of IHCTG pool compared to Huh7 cells. It is also plausible, that given the phenotype of the donors and respective intakes of alcohol, that DNL was upregulated process ^[53]^ and the PHH cells had a higher proportion of DNL-derived palmitate in the IHCTG at seeding.

When glucose is consumed *in vivo* in humans, there is a corresponding insulin response which helps co-ordinate the utilisation of glucose to minimise fluctuations in glycaemia by providing a signal for hepatic glycogen deposition, and upregulating the hepatic DNL pathway ^[54]^. We found glycogen deposition to be similar between both cell types and across all treatments, which may in part be explain by the fact, we did not add insulin into the media and therefore did not upregulate glycogen deposition; it would be of interest to see if this pathway could be upregulated in future experiments. For Huh7 cells, the lack of excess substrate, due to being cultured in lower amounts of glucose than PHH, may have contributed to the lower levels of intracellular glycogen. Alternatively, any excess glucose may have been partitioned toward glycolysis as media lactate levels were notably higher in Huh7 compared to PHH cells under all treatments, which is likely due to Huh7 being a hepatoma-derived cell line and have been reported to rely on glycolysis for rapid growth ^[55]^.

The divergence in the media concentrations of lactate and 3OHB as markers of FA and glucose oxidation by PHH and Huh7 cells, was one of the more notable differences between the cell types. This distinction may be particularly relevant for modelling MASLD where altered hepatic substrate oxidation and ketogenesis are key early metabolic adaptations. In line with our current observation of a significant decrease in media lactate with the addition of FAs, we have previously found in Huh7 cells that media lactate is significantly lower when cells were cultured in 800µM FA and 11mM glucose + 5.5mM fructose when compared to cells cultured in 200µM FA and 11mM glucose + 5.5mM fructose ^[21]^. Thus, it would be of interest to culture Huh7 longer in lower glucose and higher FA to see if they metabolically shifted away from glycolysis toward greater FA oxidation. The ketone body, 3OHB, is often used as a surrogate marker of hepatocellular FA oxidation and we found that media concentrations were approximately 3-fold lower in Huh7 compared to PHH cells, with no differences being noted across treatments. We have previously found media 3OHB to increase when Huh7 cells were cultured in 800µM FA and 11mM glucose + 5.5mM fructose compared to cells cultured in 200µM FA and 11mM glucose + 5.5mM fructose ^[21]^. The higher media 3OHB in PHH may in part explain the lower magnitude of increase in PHH IHCTG compared to Huh7 with FA treatment, with some of the excess FA being disposed via oxidation. Although we did not measure complete mitochondrial oxidation, we have previously found Huh7 cells cultured in 800µM FA and 5.5mM glucose for 7 days had higher FA oxidation compared to cells cultured in 200µM and 5.5mM glucose ^[20]^. The availability of FA to enter oxidation pathways may be influenced by how rapidly LDs turnover and FA are made available; it remains to be elucidated if differences in LD TG turnover exist between FA treatments and cell types.

Transcriptomic analysis revealed both overlap and divergence in responses of PHH and Huh7 cells treated with FA. In Huh7 cells, the dominant source of transcriptional variance was FA-treatment, where in PHH cells transcriptomic variability was driven by primarily by donor-specific differences, suggesting that the genotype and baseline phenotype of these cells have a far greater impact on transcription than treatment with FAs. Transcriptomic analysis suggested that Huh7 cells become physiologically adapted to these FA mixtures after chronic exposure with a shift towards oxidative phosphorylation and FA metabolism with reduced stress and inflammatory responses. However, this muted inflammatory response in Huh7 cells likely reflects their transformed origin and reduced competence for stress and immune signalling, meaning PHH remain essential for modelling inflammatory aspects of MASLD progression. In contrast, PHH cells displayed activation of stress-related pathways with FA treatment, together with suppression of lipid metabolic pathways. This divergence highlights a fundamental biological difference between Huh7 and PHH cells in modelling aspects of FA induced hepatocellular stress.

The absence of inflammatory pathway activation in Huh7 cells may reflect their transformed origin and altered stress-response signalling, whereas the inflammatory transcriptional response observed in PHH cells is likely influenced by higher baseline intracellular lipid content and reduced cellular viability following cryopreservation. These factors may also constrain the dynamic range of transcriptional responses in PHH and contribute to the observed differences between OPLA- and POLA-treated conditions across cell types. Responses may also reflect, in part, culture media, duration of FA exposure, along with how long IHCTG had been accumulating, as PHH cells have been exposed to more than FAs and likely further along the liver disease pathology pathway.

Cells used in *in vitro* experiments should have high viability, indicating they are live, healthy, and functional, so the effect of a treatment or specific conditions can be robustly assessed. Typically, viability assays assess membrane integrity, metabolic activity, or enzymatic function. We assessed cell viability in two ways, using trypan blue prior to seeding, where cells with compromised membranes take up the dye, and by measuring ATP levels after cells had treatment media for 4-7 days, as a proxy for cellular energy state. For both assays, PHH demonstrated lower viability compared with Huh7 cells. These observations are consistent with prior reports that highlight the reduced proliferative capacity and metabolic stability of PHH post-cryopreservation ^[56]^, along with the variability in viability noted in the certificate of analysis that is provided with the PHH when purchased.

### Limitations of this study

Our study is not without limitations. We used trypan blue staining at seeding to determine cell viability and found it to be lower in PHH cells. As PHH were cultured in collagen-based adherence methods, non-viable cells could not be removed and may therefore have contributed to downstream measurements potentially leading an underestimation in cell viability. Culture duration differed between the two cell types, with PHHs, due to lower viability, being maintained for only 4 days (media changes every 2 days), thus, limiting our ability to explore chronic FA exposure and Huh7 cells were cultured for 7 days (media changes every 2 days). In addition, basal media composition differed between models, including glucose concentration, which may have influenced substrates utilisation and metabolic fluxes. Furthermore, donor metabolic history prior to hepatocytes isolation is not fully known, which may contribute to baseline lipid variability and transcriptional differences observed in PHH. Therefore, in future studies, where possible, we would culture both cell types for the same duration, under, where possible similar substrate conditions. We used 2-dimensional culturing and it would be of interest to use our media with a physiologically relevant three-dimensional culture system as such an approach may better capture haptic zonation, oxygen gradients and inter cellular interaction that influence hepatocytes metabolism *in-vivo*.

## Conclusion

Overall, our data indicate that while immortalized hepatic cell lines are often considered to be a poor surrogate for PHH metabolism, Huh7 chronically cultured in physiologically relevant media, are metabolically adaptable and show comparable responses that overlap with PHH in selected lipid and metabolic readouts. In particular, similarities were observed in FA uptake, IHCTG composition and LD characteristics where cells were exposed to physiological levels and composition of FA. These shared features coexist with clear biological differences between the two models. PHH displayed higher baseline IHCTG content and greater donor dependent variability, where Huh7 showed more uniform metabolic and transcriptomic adaptation to chronic FA exposure. Taken together our findings suggest there as Huh7 are metabolically adaptable and with further optimization, they could better recapitulate PHH metabolism, which is dependent on the phenotype and genotype of the donor, thus making Huh7 a practical and scalable workhorse model to investigate relevant pathways that may underpin the development of MASLD.

## Author contributions

LH, EC, FW, SRN and FS designed the research, EC, FW, KS, SRN, and KD conducted the research, EC, FW, KS and KD analysed the data and performed statistical analysis and drafted the manuscript. KD and LH have primary responsibility for final content. All authors have read and approved the final manuscript.

## REFERENCES

[1] Rinella ME, Lazarus JV, Ratziu V, et al. A multisociety Delphi consensus statement on new fatty liver disease nomenclature. Hepatology. 2023;78:1966–1986.

[2] Di Mauro S, Scamporrino A, Filippello A, et al. Clinical and Molecular Biomarkers for Diagnosis and Staging of NAFLD. Int J Mol Sci. 2021;22.

[3] Younossi ZM, Golabi P, Paik JM, et al. The global epidemiology of nonalcoholic fatty liver disease (NAFLD) and nonalcoholic steatohepatitis (NASH): a systematic review. Hepatology. 2023;77:1335–1347.

[4] Jamialahmadi O, Tavaglione F, Rawshani A, et al. Fatty liver disease, heart rate and cardiac remodelling: Evidence from the UK Biobank. Liver Int. 2023;43:1247–1255.

[5] Diraison F, Beylot M. Role of human liver lipogenesis and reesterification in triglycerides secretion and in FFA reesterification. Am J Physiol. 1998;274:E321–327.

[6] Sidossis LS, Mittendorfer B, Walser E, et al. Hyperglycemia-induced inhibition of splanchnic fatty acid oxidation increases hepatic triacylglycerol secretion. Am J Physiol. 1998;275:E798–805.

[7] Babin PJ, Gibbons GF. The evolution of plasma cholesterol: direct utility or a “spandrel” of hepatic lipid metabolism? Prog Lipid Res. 2009;48:73–91.

[8] Hodson L, Frayn KN. Hepatic fatty acid partitioning. Curr Opin Lipidol. 2011;22:216–224.

[9] Lau JK, Zhang X, Yu J. Animal models of non-alcoholic fatty liver disease: current perspectives and recent advances. J Pathol. 2017;241:36–44.

[10] Li JZ, Huang Y, Karaman R, et al. Chronic overexpression of PNPLA3I148M in mouse liver causes hepatic steatosis. J Clin Invest. 2012;122:4130–4144.

[11] Ding Y, Dai X, Bao M, et al. Hepatic transcriptome signatures in mice and humans with nonalcoholic fatty liver disease. Animal Model Exp Med. 2023;6:317–328.

[12] Green CJ, Pramfalk C, Morten KJ, et al. From whole body to cellular models of hepatic triglyceride metabolism: man has got to know his limitations. Am J Physiol Endocrinol Metab. 2015;308:E1–20.

[13] Kwon Y, Gottmann P, Wang S, et al. Induction of steatosis in primary human hepatocytes recapitulates key pathophysiological aspects of metabolic dysfunction-associated steatotic liver disease. J Hepatol. 2024.

[14] Huggett ZJ, Smith A, De Vivo N, et al. A Comparison of Primary Human Hepatocytes and Hepatoma Cell Lines to Model the Effects of Fatty Acids, Fructose and Glucose on Liver Cell Lipid Accumulation. Nutrients. 2022;15.

[15] Godoy P, Hewitt NJ, Albrecht U, et al. Recent advances in 2D and 3D in vitro systems using primary hepatocytes, alternative hepatocyte sources and non-parenchymal liver cells and their use in investigating mechanisms of hepatotoxicity, cell signaling and ADME. Arch Toxicol. 2013;87:1315–1530.

[16] Gunn PJ, Green CJ, Pramfalk C, et al. In vitro cellular models of human hepatic fatty acid metabolism: differences between Huh7 and HepG2 cell lines in human and fetal bovine culturing serum. Physiol Rep. 2017;5.

[17] Pramfalk C, Larsson L, Hardfeldt J, et al. Culturing of HepG2 cells with human serum improve their functionality and suitability in studies of lipid metabolism. Biochim Biophys Acta. 2016;1861:51–59.

[18] Steenbergen R, Oti M, Ter Horst R, et al. Establishing normal metabolism and differentiation in hepatocellular carcinoma cells by culturing in adult human serum. Sci Rep. 2018;8:11685.

[19] Steenbergen RH, Joyce MA, Thomas BS, et al. Human serum leads to differentiation of human hepatoma cells, restoration of very-low-density lipoprotein secretion, and a 1000-fold increase in HCV Japanese fulminant hepatitis type 1 titers. Hepatology. 2013;58:1907–1917.

[20] Nagarajan SR, Cross E, Johnson E, et al. Determining the temporal, dose, and composition effects of nutritional substrates in an in vitro model of intrahepatocellular triglyceride accumulation. Physiol Rep. 2022;10:e15463.

[21] Gunn PJ, Pramfalk C, Millar V, et al. Modifying nutritional substrates induces macrovesicular lipid droplet accumulation and metabolic alterations in a cellular model of hepatic steatosis. Physiol Rep. 2020;8:e14482.

[22] Robinson MD, Oshlack A. A scaling normalization method for differential expression analysis of RNA-seq data. Genome Biol. 2010;11:R25.

[23] Robinson MD, McCarthy DJ, Smyth GK. edgeR: a Bioconductor package for differential expression analysis of digital gene expression data. Bioinformatics. 2010;26:139–140.

[24] Ritchie ME, Phipson B, Wu D, et al. limma powers differential expression analyses for RNA-sequencing and microarray studies. Nucleic Acids Res. 2015;43:e47.

[25] Liberzon A, Birger C, Thorvaldsdottir H, et al. The Molecular Signatures Database (MSigDB) hallmark gene set collection. Cell Syst. 2015;1:417–425.

[26] Smith K, Dennis K, Hodson L. The ins and outs of liver fat metabolism: The effect of phenotype and diet on risk of intrahepatic triglyceride accumulation. Exp Physiol. 2025;110:936–948.

[27] Luukkonen PK, Sadevirta S, Zhou Y, et al. Saturated Fat Is More Metabolically Harmful for the Human Liver Than Unsaturated Fat or Simple Sugars. Diabetes Care. 2018;41:1732–1739.

[28] Parry SA, Rosqvist F, Mozes FE, et al. Intrahepatic Fat and Postprandial Glycemia Increase After Consumption of a Diet Enriched in Saturated Fat Compared With Free Sugars. Diabetes Care. 2020;43:1134–1141.

[29] Rosqvist F, Iggman D, Kullberg J, et al. Overfeeding polyunsaturated and saturated fat causes distinct effects on liver and visceral fat accumulation in humans. Diabetes. 2014;63:2356–2368.

[30] Rosqvist F, Kullberg J, Stahlman M, et al. Overeating Saturated Fat Promotes Fatty Liver and Ceramides Compared With Polyunsaturated Fat: A Randomized Trial. J Clin Endocrinol Metab. 2019;104:6207–6219.

[31] Breher-Esch S, Sahini N, Trincone A, et al. Genomics of lipid-laden human hepatocyte cultures enables drug target screening for the treatment of non-alcoholic fatty liver disease. BMC Med Genomics. 2018;11:111.

[32] Liu X, Green RM. Endoplasmic reticulum stress and liver diseases. Liver Res. 2019;3:55–64.

[33] Maruyama H, Takahashi M, Sekimoto T, et al. Linoleate appears to protect against palmitate-induced inflammation in Huh7 cells. Lipids Health Dis. 2014;13:78.

[34] Moliterni C, Vari F, Schifano E, et al. Lipotoxicity of palmitic acid is associated with DGAT1 downregulation and abolished by PPARalpha activation in liver cells. J Lipid Res. 2024;65:100692.

[35] Wei Y, Wang D, Topczewski F, et al. Saturated fatty acids induce endoplasmic reticulum stress and apoptosis independently of ceramide in liver cells. Am J Physiol Endocrinol Metab. 2006;291:E275–281.

[36] de Sousa IF, Migliaccio V, Lepretti M, et al. Dose- and Time-Dependent Effects of Oleate on Mitochondrial Fusion/Fission Proteins and Cell Viability in HepG2 Cells: Comparison with Palmitate Effects. Int J Mol Sci. 2021;22.

[37] Eynaudi A, Diaz-Castro F, Borquez JC, et al. Differential Effects of Oleic and Palmitic Acids on Lipid Droplet-Mitochondria Interaction in the Hepatic Cell Line HepG2. Front Nutr. 2021;8:775382.

[38] Gomez-Lechon MJ, Donato MT, Martinez-Romero A, et al. A human hepatocellular in vitro model to investigate steatosis. Chem Biol Interact. 2007;165:106–116.

[39] Lee JY, Cho HK, Kwon YH. Palmitate induces insulin resistance without significant intracellular triglyceride accumulation in HepG2 cells. Metabolism. 2010;59:927–934.

[40] Listenberger LL, Han X, Lewis SE, et al. Triglyceride accumulation protects against fatty acid-induced lipotoxicity. Proc Natl Acad Sci U S A. 2003;100:3077–3082.

[41] Ricchi M, Odoardi MR, Carulli L, et al. Differential effect of oleic and palmitic acid on lipid accumulation and apoptosis in cultured hepatocytes. J Gastroenterol Hepatol. 2009;24:830–840.

[42] Moravcova A, Cervinkova Z, Kucera O, et al. The effect of oleic and palmitic acid on induction of steatosis and cytotoxicity on rat hepatocytes in primary culture. Physiol Res. 2015;64:S627–636.

[43] Teixeira FS, Pimentel LL, Vidigal S, et al. Differential Lipid Accumulation on HepG2 Cells Triggered by Palmitic and Linoleic Fatty Acids Exposure. Molecules. 2023;28.

[44] Hodson L, Skeaff CM, Fielding BA. Fatty acid composition of adipose tissue and blood in humans and its use as a biomarker of dietary intake. Prog Lipid Res. 2008;47:348–380.

[45] Lagziel S, Gottlieb E, Shlomi T. Mind your media. Nat Metab. 2020;2:1369–1372.

[46] Paramitha PN, Zakaria R, Maryani A, et al. Raman Study on Lipid Droplets in Hepatic Cells Co-Cultured with Fatty Acids. Int J Mol Sci. 2021;22.

[47] Scorletti E, Carr RM. A new perspective on NAFLD: Focusing on lipid droplets. J Hepatol. 2022;76:934–945.

[48] Solinas G, Boren J, Dulloo AG. De novo lipogenesis in metabolic homeostasis: More friend than foe? Mol Metab. 2015;4:367–377.

[49] Lambert JE, Ramos-Roman MA, Valdez MJ, et al. Weight loss in MASLD restores the balance of liver fatty acid sources. J Clin Invest. 2025;135.

[50] Luukkonen PK, Porthan K, Ahlholm N, et al. The PNPLA3 I148M variant increases ketogenesis and decreases hepatic de novo lipogenesis and mitochondrial function in humans. Cell Metab. 2023;35:1887–1896 e1885.

[51] Pramfalk C, Pavlides M, Banerjee R, et al. Fasting Plasma Insulin Concentrations Are Associated With Changes in Hepatic Fatty Acid Synthesis and Partitioning Prior to Changes in Liver Fat Content in Healthy Adults. Diabetes. 2016;65:1858–1867.

[52] Smith GI, Shankaran M, Yoshino M, et al. Insulin resistance drives hepatic de novo lipogenesis in nonalcoholic fatty liver disease. J Clin Invest. 2020;130:1453–1460.

[53] Siler SQ, Neese RA, Hellerstein MK. De novo lipogenesis, lipid kinetics, and whole-body lipid balances in humans after acute alcohol consumption. Am J Clin Nutr. 1999;70:928–936.

[54] Cross E, Dearlove DJ, Hodson L. Nutritional regulation of hepatic de novo lipogenesis in humans. Curr Opin Clin Nutr Metab Care. 2023;26:65–71.

[55] Kiss I, Neuwert N, Oberle R, et al. Hepatic Lipoprotein Metabolism: Current and Future In Vitro Cell-Based Systems. Biomolecules. 2025;15.

[56] Sun Z, Yuan X, Wu J, et al. Hepatocyte transplantation: The progress and the challenges. Hepatol Commun. 2023;7.

