## Supplemental Table 1 for "Physiologically relevant media are associated with overlapping metabolic responses in primary human hepatocytes and Huh7 cells"

**Supplementary Table 1:** Primary Human Hepatocyte Donor Characteristics

| **Sex** | **Race** | **Age (y)** | **BMI**  **(kg/m^2^)** | **Tobacco history** | **Alcohol history** | **Drug history** | **Cause of death** | **Viable cells/Vial*** |
| --- | --- | --- | --- | --- | --- | --- | --- | --- |
| F | AA | 31 | 18.9 | Y | Y | Marijuana | Asphyxiation | 9.0 x10^6^ |
| F | AA | 65 | 39.6 | Y | Y | Bydureon, Amlodipine, Carvedilol, Losartan, Hydrochlorothiazide, Clonidine, Atorvastatin, Aspirin, Clopidogrel, Cholecalciferol | Unknown | 9.7 x10^6^ |
| F | C | 55 | 22.8 | 2-3 cpd for 2mths | 4 dpd for 32 y | Cannabis daily for 5y | Brain tumour | 4.9 x10^6^ |
| M | C | 46 | 32.1 | 1 ppd 34y | 2-4 dpm | Cannabis for 30 years, cocaine for 1y | Asphyxiation | 7.42 x10^6^ |
| M | A | 47 | 23.9 | 1 ppd 20y | 6 beers dpd for 20 y | Cannabis daily for 20y | Anoxia /Stroke | 11.34 x10^6^ |

Abbreviations: F, Female; M, Male; AA, African American; C, Caucasian, A, Asian; Y, yes; cpd, cigarettes per day; mth, months; ppd, packet per day;y, years; dpd, drinks per day; dpm, drinks per month. *post thaw results
