## Supplementary figures and images for "Physiologically relevant media are associated with overlapping metabolic responses in primary human hepatocytes and Huh7 cells"

### Supplemental Figure 1

Supplementary Figure 1.

A.

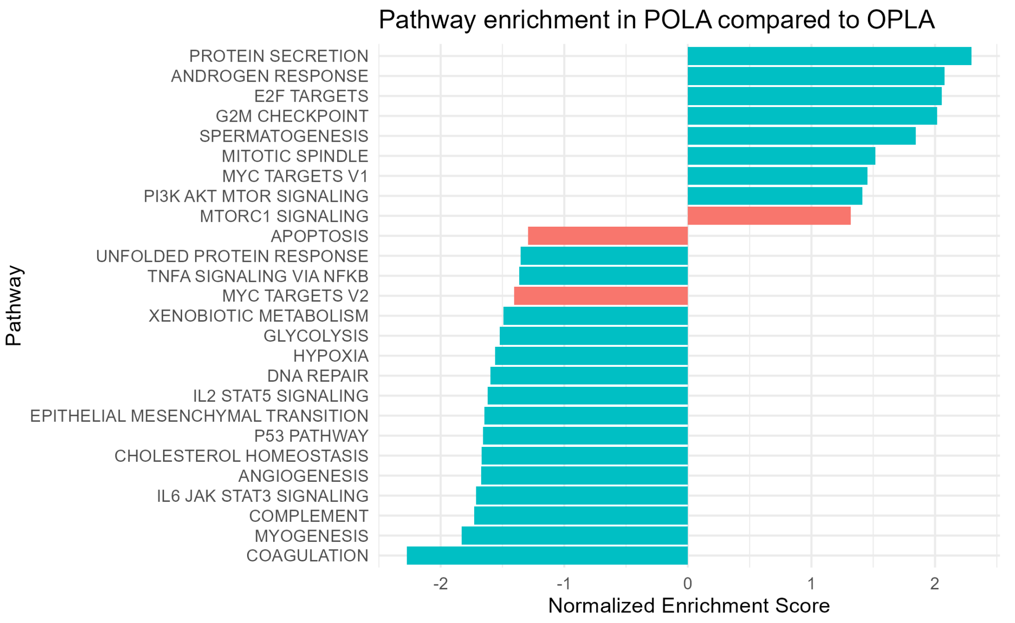


B.

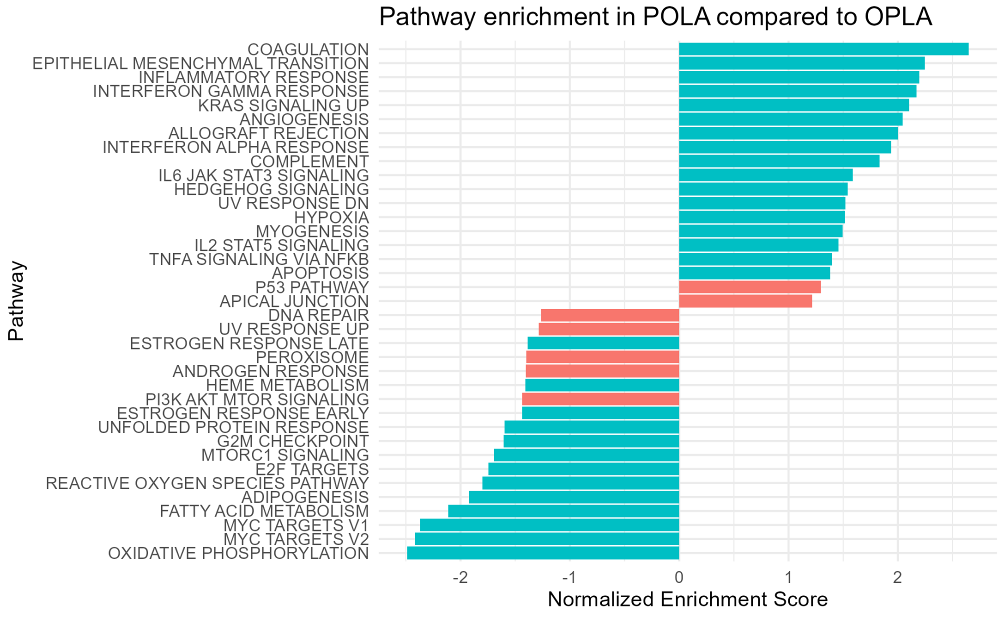
